## Supplementary material for "Picoplankton nitrogen guilds in the tropical and subtropical oceans: from the surface to the deep": Supplementary11.pdf

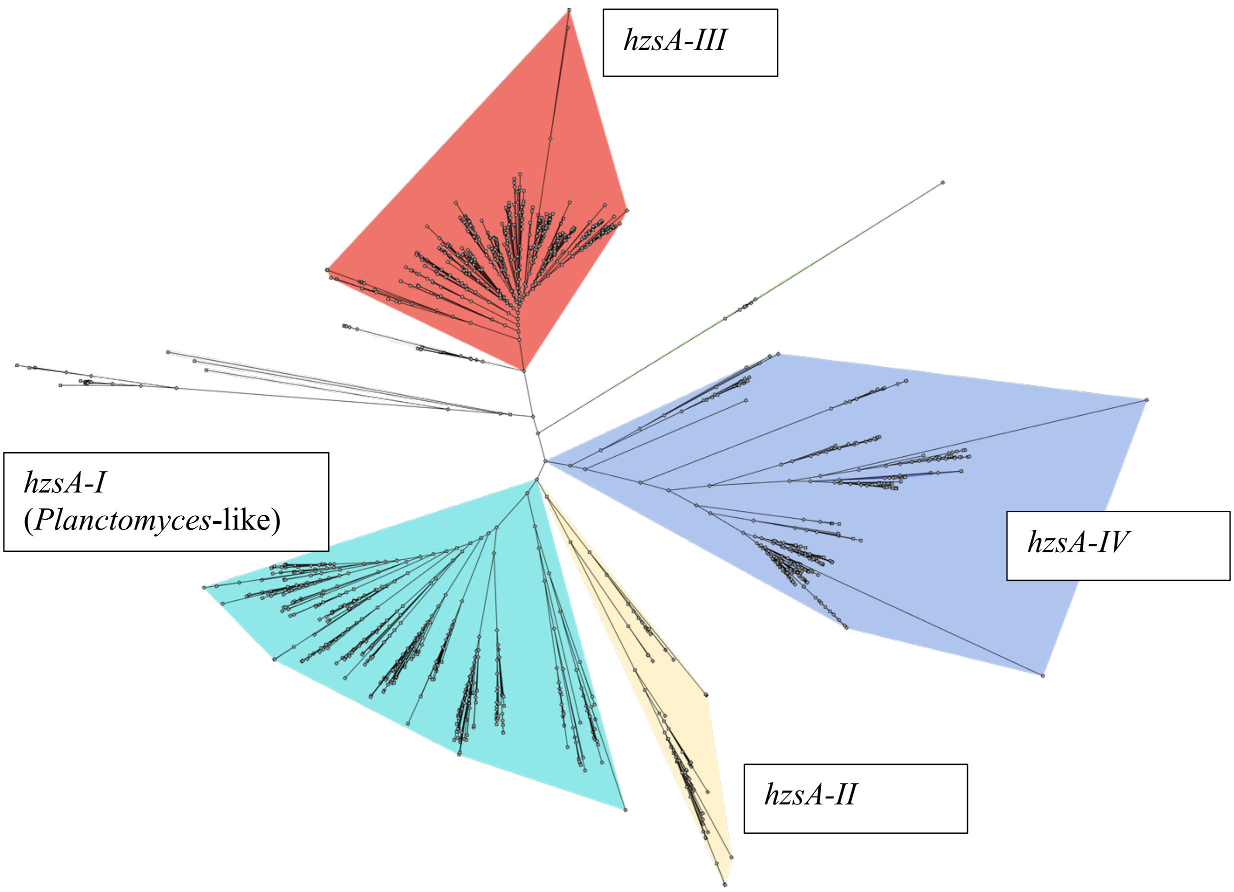


**Figure S1** – Reference tree for the hzsA gene. The tree was built with sequences from data bases and showed four main clusters (I to IV). Cluster I contains most *Planctomycetes* sequences and a few from *Verrucomicrobia*. The other three clusters contain sequences from microorganisms not know to carry out annamox. Then, metagenomic reads were placed in the tree. Sequences from the OMZ were placed in cluster I as expected. Sequences from station SAtl were surprisingly placed in cluster III and belonged to *Alcanivorax* relatives. This shows that our method is able to identify paralogs that would be erroneously counted by conventional annotation. Perhaps the cluster III proteins carry out some function related to the hydrocarbon degradation processes for which *Alcanivorax* is known

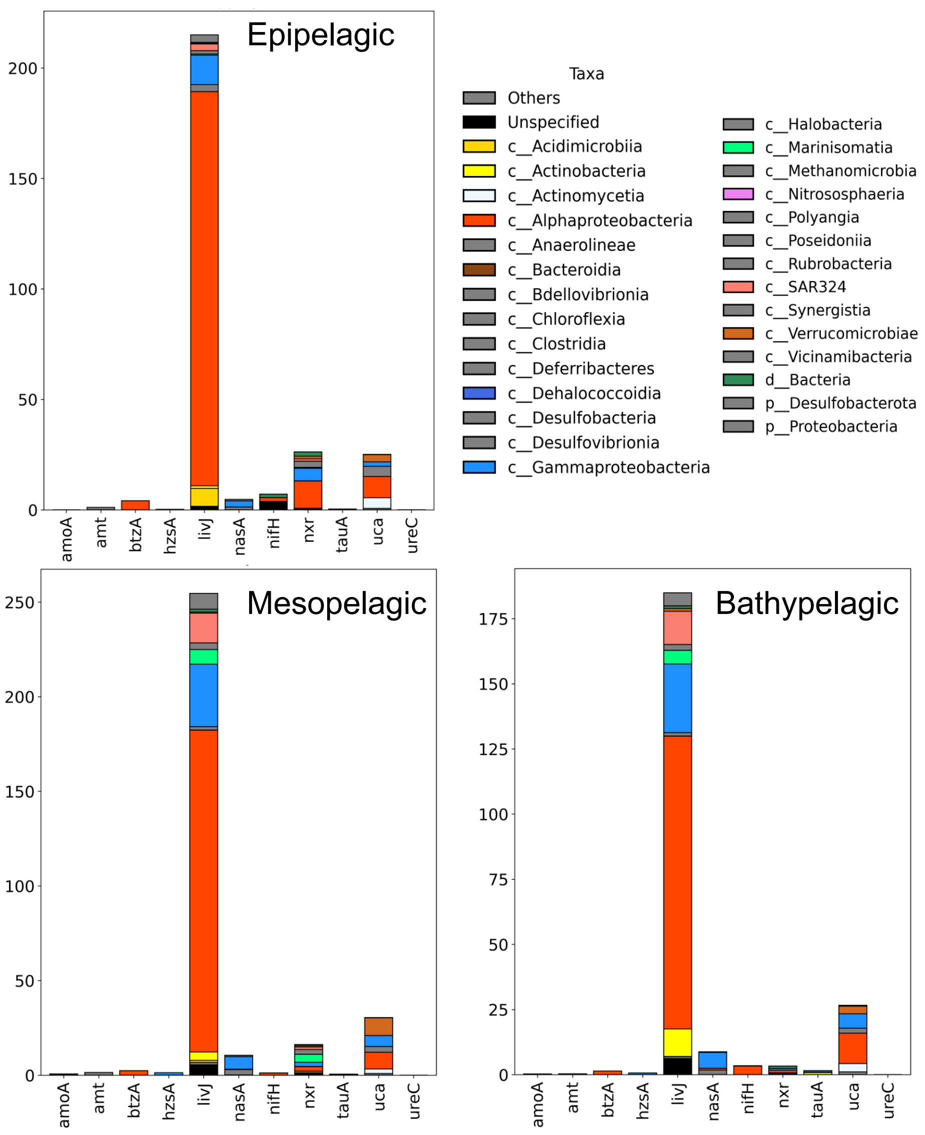


**Figure S2** – Discarded reads (FPM) after the curated annotation for the most abundant genes. The livJ gene, one of the most abundant, was also the one with the most reads rejected. However, the number of discarded reads did not depend only on the abundance of the gene. For example, amt (one with the most reads) lost almost no reads, while nifH (with very low number of reads) lost most of them. Rather, this depends on the closeness of the gene sequences to those of its paralogs.

N uptake

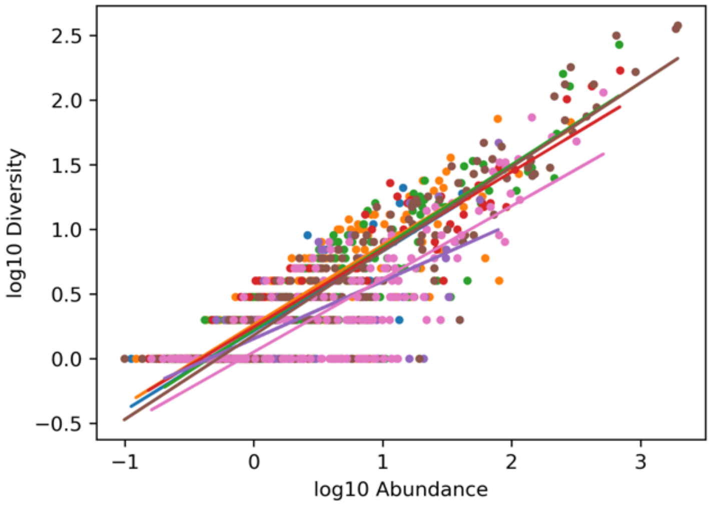


N redox

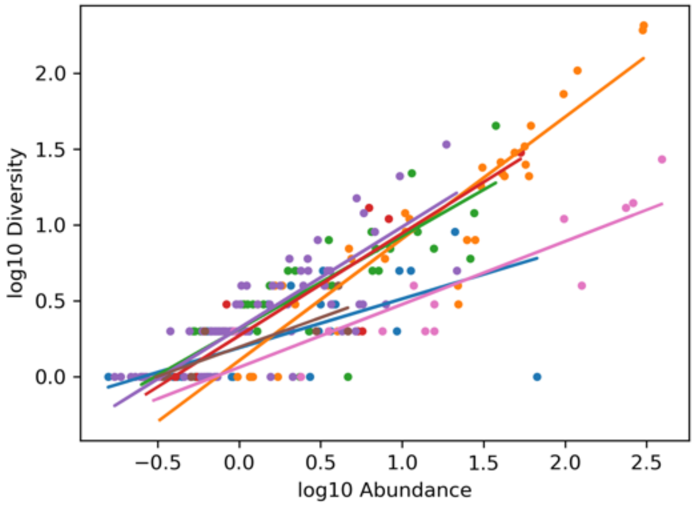


**Figure S3** – Calculation for expected diversity (∂exp) from abundance (A) for different genes. The observed diversity (∂obs) was plotted against the abundance in a log-log scale. The slope of the regression allows calculation of the diversity expected from any given abundance. Then, the quotient ∂obs / ∂exp provides de delta value.

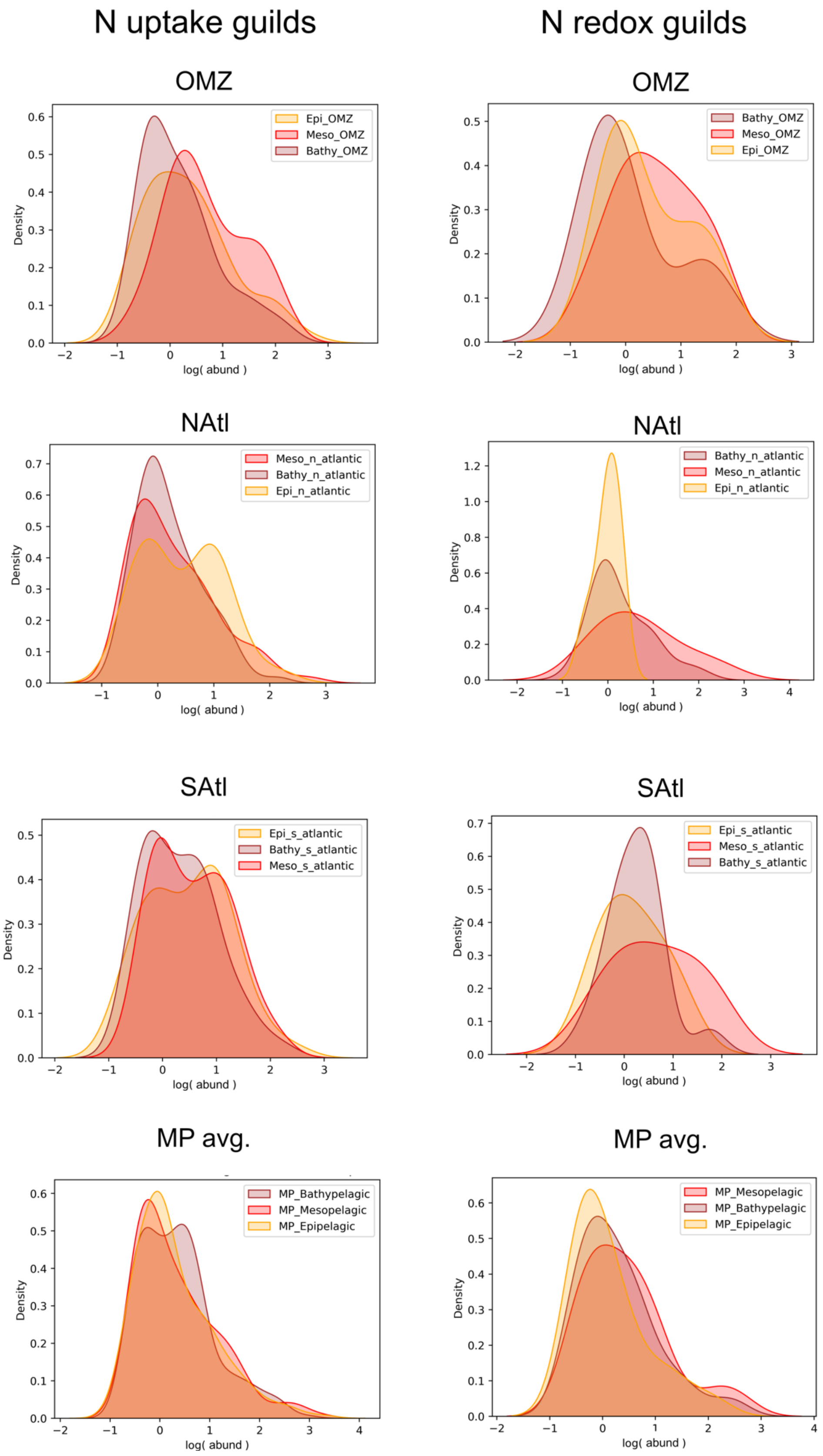


**Figure S4** – Density functions for abundance values for the different depths and stations analyzed. These graphs are equivalent to those in figure 5 but for abundance instead of delta values. Abundance distributions are closer among N uptake and N redox genes than those for diversity. Thus, abundance alone misses relevant information. We had more data for N uptake than for N redox functions. This made the inference for the former more robust, albeit r-squared values were similar (full list provided in Table S3).

### Supplementary Files

**File S1:** All reference trees and data needed for the various metagenomic placements.

### Supplementary Tables

**Table S1:** List of the studied genes encoding nitrogen-cycling proteins. The corresponding genes were searched in each sample using the indicated protein models.

**Table S2:** All the metagenomic queries (functionally annotated with abundances, filtered and unfiltered) used in this work.

**Table S3:** All the statistical tests performed regarding diversification of protein sequences.
